# Prior exposure to speech rapidly modulates cortical processing of high-level linguistic structure

**DOI:** 10.1101/2022.01.25.477669

**Authors:** Qingqing Meng, Yiwen Li Hegner, Iain Giblin, Catherine McMahon, Blake W Johnson

**Affiliations:** The HEARing CRC, Audiology, Hearing and Speech Sciences, University of Melbourne, Melbourne, Victoria, Australia; Macquarie University, Department of Cognitive Science, Sydney, New South Wales, Australia; Macquarie University, Department of Linguistics, Sydney, New South Wales, Australia; H:EAR Centre, Macquarie University, New South Wales, Australia; MEG-Center, University of Tübingen, Tübingen, Germany; National Acoustic Laboratories, Australian Hearing Hub, Sydney, New South Wales, Australia

**Keywords:** Brain imaging, magnetoencephalography, prior knowledge, speech intelligibility

## Abstract

Neural activity has been shown to track hierarchical linguistic units in connected speech and these responses can be directly modulated by changes in speech intelligibility caused by spectral degradation. In the current study, we manipulate prior knowledge to increase the intelligibility of physically identical speech sentences and test the hypothesis that the tracking responses can be enhanced by this intelligibility improvement. Cortical magnetoencephalography (MEG) responses to intelligible speech followed by either the same (matched) or different (unmatched) unintelligible speech were measured in twenty-three normal hearing participants. Driven by prior knowledge, cortical coherence to “abstract” linguistic units with no accompanying acoustic cues (phrases and sentences) was enhanced relative to the unmatched condition, and was lateralized to the left hemisphere. In contrast, cortical responses coherent to word units, aligned with acoustic onsets, were bilateral and insensitive to contextual information changes. No such coherence changes were observed when prior experience was not available (unintelligible speech before intelligible speech). This dissociation suggests that cerebral responses to linguistic information are directly affected by intelligibility, which in turn are powerfully shaped by physical cues in speech. These results provide an objective and sensitive neural index of speech intelligibility, and explain why previous studies have reported no effect of prior knowledge on cortical entrainment.

## 1. Introduction

A dramatic enhancement in the perceived intelligibility of distorted speech signal can be achieved by providing prior information on the content of the signal (Jacoby, Allan, Collins, & Larwill, 1988; Remez, Rubin, Pisoni, & Carrell, 1981). This top-down perceptual change, referred to as perceptual “pop-out” (Davis, Johnsrude, Hervais-Adelman, Taylor, & McGettigan, 2005), is invoked rapidly and reliably with immediate prior exposure to a clear speech signal.

Several neuroimaging studies have examined changes in brain activity associated with this perceptual “pop-out” effect. In a series of functional magnetic resonance imaging (fMRI) studies, decreased activations have been reported in the left superior temporal lobe and right lateral Heschl’s gyrus (HG) (Liebenthal, Binder, Piorkowski, & Remez, 2003) while increased activations have been identified in the right anterior superior temporal sulcus and a set of regions of the bilateral middle and inferior temporal gyri (Giraud et al., 2004), the posterior part of the left superior temporal gyrus (STG) extending along the superior temporal sulcus (Dehaene-Lambertz et al., 2005) and the planum temporale and planum polare, and the superior/middle temporal gyri extending into inferior parietal and frontal cortices (Tuennerhoff & Noppeney, 2016). Neurophysiological measures with EEG (electroencephalography) and MEG (magnetoencephalography) have also been employed to investigate the temporal profiles of auditory neural responses. An electrophysiological mismatch response (MMR) has been reported to occur earlier and more asymmetrically for a phonemic change than for an equivalent acoustic change (Dehaene-Lambertz et al., 2005). Prior knowledge-induced EEG enhancement and concurrent MEG reduction have been localized to the inferior frontal gyrus (IFG) and left STG (Sohoglu, Peelle, Carlyon, & Davis, 2012; Sohoglu & Davis, 2016), with a temporal ordering such that the activity in IFG was modulated before the activity in lower-level sensory regions of the left STG. Using a pop-out paradigm, a positive correlation between delta band entrainment to phoneme-level features and perceived speech intelligibility has been reported (Liberto, Crosse, & Lalor, 2018). Furthermore, a significant effect of prior knowledge on cortical entrainment to the temporal envelope speech has been reported in distinct time windows in the left IFG and HG (Di Liberto, Lalor, & Millman, 2018).

A recent invasive electrocorticography (ECoG) study has quantified changes in the spectrotemporal tuning of ensemble neuronal activity with recordings obtained directly from human auditory cortex (Holdgraf et al., 2016). This tuning or feature representation, described as the neurons’ spectrotemporal receptive fields (STRFs), has conventionally been examined in animal models using single-unit recordings at different levels of the auditory pathway (Miller, Escabí, Read, & Schreiner, 2002; Woolley, Fremouw, Hsu, & Theunissen, 2005). Based on ensemble spectrotemporal receptive fields (eSTRFs), Holdgraf and colleagues demonstrated a rapid automatic change of speech feature encoding in human auditory cortex, induced by prior experience of intact speech before subsequent presentations of degraded speech. This tuning shift has been suggested to facilitate extraction of speech related features in stimuli and provide the physiological basis for the experience-enhanced speech pop-out phenomenon (Holdgraf et al., 2016).

In the context of human speech perception, neurophysiological studies have shown that the auditory cortex tracks the dynamics of speech envelope, approximately at the syllabic rate (Ahissar et al., 2001; Ding & Simon, 2012; Kayser, Ince, Gross, & Kayser, 2015; Lakatos et al., 2005; Luo & Poeppel, 2007; Rimmele, Zion Golumbic, Schröger, & Poeppel, 2015).

Psychoacoustic studies have shown that the slowly varying temporal envelope of speech signal contains major acoustic cues that are important for speech intelligibility (Drullman, Festen, & Plomp, 1994; Shannon, Zeng, Kamath, Wygonski, & Ekelid, 1995; Smith, Delgutte, & Oxenham, 2002). This neural tracking activity, often referred to as “cortical entrainment” has been argued to be necessary for speech comprehension. However, its functional role remains controversial (Ding & Simon, 2014; Peelle & Davis, 2012; Zoefel & VanRullen, 2015). Some authors believe that cortical synchronization with the low-frequency speech envelope actively constrains the transfer of information from sensory to higher-order brain regions and this synchronization with the speech envelope is essential for speech comprehension (Ding, Chatterjee, & Simon, 2014; Peelle, Gross, & Davis, 2013; Zion Golumbic et al., 2013). Others have argued that the role of phase-locking brain responses may be restricted to encoding acoustic cues at the syllabic rhythm in speech (Nourski et al., 2009; Howard & Poeppel, 2010; Doelling, Arnal, Ghitza, & Poeppel, 2014), and that the cortical responses are mainly driven by the physical properties of the acoustic input. Due to the concomitant changes in intelligibility and acoustics properties in speech stimuli, there has been a continuing debate about whether the brain envelope-following response mainly reflects processing of linguistic or acoustic information in speech.

By manipulating prior knowledge of spoken sentences, the effects of perceived intelligibility on cortical entrainment can be easily isolated from any acoustical changes in speech stimuli. However, counter to expectations and the behaviourally robust perceptual enhancement in speech intelligibility, none of the studies employing the pop-out paradigm have reported any significant effects of prior knowledge on brain activities phase-locked to the temporal envelope of speech (Holdgraf et al., 2016; Liberto et al., 2018; Millman, Johnson, & Prendergast, 2014). One study has reported significant cortical entrainment enhancement in the delta band (1-4 Hz) induced by perceptual pop-out, however rather than sustained throughout the duration of speech utterance it only emerged within overlapped time windows up to 400ms (Di Liberto et al., 2018).

An important methodological advance has been provided by MEG work demonstrating that activity from auditory cortex can track *abstract* linguistic units, i.e., linguistic units that are embedded in connected speech but have no physical presence in the acoustic properties of the signal (Ding, Melloni, Zhang, Tian, & Poeppel, 2016). When short sentences constructed with the same syntactic structure were presented in an isochronous manner, concurrent cortical tracking activity to syllable/word, phrase and sentence level linguistic units from participants was found. Importantly, this neural tracking activity of larger linguistic structure at phrase and sentence level is unambiguously dissociated from encoding of acoustic cues to these units, because there are no physical phrase or sentence boundaries in the isochronous speech signal. The authors argued that an internal, grammar-based construction process must have been implemented (Ding et al., 2016).

Our recent MEG study has demonstrated the effect of spectral degradation modulated speech intelligibility on these cortical tracking responses and investigated the underlying neural sources of the concurrent tracking responses (Meng, Li Hegner, Giblin, McMahon, & Johnson, 2021). Results of this study showed that cortical entrainment – the coherence between brain activities and “abstract” linguistic units with no accompanying acoustic cues (phrases and sentences) – was reduced parametrically as a function of reduced intelligibility. In contrast, brain responses coherent to words/syllables that were accompanied by acoustic onsets were insensitive to intelligibility changes. Beam-forming source localization analysis further demonstrated that the intelligibility-modulated brain tracking activities were lateralized to the left hemisphere while the intelligibility-insensitive word/syllable level tracking responses were bilateral. Importantly, these results indicated that brain responses are relatively insensitive to changes in intelligibility when linguistic and acoustic temporal regularities are mixed up together. This confound between acoustic and linguistic cues is inevitable in naturalistic speech and it may account for the mixed results reported in neuroimaging studies that employed naturalistic sentences for experimental stimuli.

Unlike previous studies, the experimental paradigm of Ding et al. (2016) provides a capable tool that unambiguously separate linguistic and acoustic cues in the speech stream and enables the assessment of intelligibility effects on neural responses at distinct timescales (syllable, phrase and sentence). Based on this paradigm and the direct modulation from intelligibility changes established in the work by Meng et al. (2021), we hypothesised that neural responses to *abstract* linguistic structures should be directly enhanced by experience-facilitated increases in speech intelligibility; while responses to linguistic regularities that are accompanied by physical cues in the speech waveform should be minimally affected by this intelligibility manipulation. Noise-vocoding was used to render speech unintelligible while maintaining its temporal envelope (Shannon et al., 1995). Enhancement in speech intelligibility were achieved via a rapid perceptual learning process (Davis et al., 2005), which changes the perceptual experience of spectrally-degraded speech sentences from unintelligible to intelligible by pre-exposing listeners to matched clear speech. In this way, all the physical properties of the speech stimuli were identical before and after the change of intelligibility was introduced.

In addition to the reported *cerebral lateralization* of brain tracking responses to lower and more abstract levels of linguistic content in our previous MEG study (Meng et al., 2021), we also wished to characterize and contrast the underlying neural sources of the tracking responses facilitated by the perceptual pop-out effect. We specifically predicted that the hypothesised enhancement in tracking responses at the sentence- and phrase-level, driven by prior knowledge on the acoustic and linguistic information, would be more left-lateralised while the largely unchanged word-level responses remain bilateral.

## 2. Materials & Methods

### 2.1. Participants

23 native speakers of English aged between 18 to 39 years old (mean 26 years old; 15 females) participated in this experiment. All participants were right-handed, with normal hearing and without any history of neurological, psychiatric, or developmental disorders (self-reported). Written informed consent was obtained from all participants under the process approved by the Human Subjects Ethics Committee of Macquarie University.

### 2.2. Stimuli

The speech materials were synthesized using the MacinTalk text to speech synthesizer (male voice Alex, 360 words per minute, Mac OS X 10.13.4). In total, 180 four-syllable (a monosyllabic word for each syllable) English sentences were generated to form a sentence list (**Supplementary Material**). All sentences in the list followed the same syntactic structures: adjective/pronoun + noun + verb + noun. Each syllable was synthesized independently, and all the synthesized syllables (200 – 376 ms in duration) were adjusted to 320 ms by truncation or padding silence at the end. The offset of each syllable was smoothed by a 25-ms cosine window.

From the 180 sentences in the total pool, 60 (first set) were randomly selected to be presented in the unprocessed form (“natural speech”). A second set of 60 sentences were randomly selected from the remaining 120 sentences for “8 channel noise vocoding”, and the remaining set (third set) of 60 sentences were used for both “natural speech” and “8 channel noise vocoding”.

With half “natural speech” and half “8 channel noise vocoding”, 12 sentences were presented in each trial. To avoid any potential artefact from the switching of acoustic conditions at the individual sentence level, every two sentences of the same type were grouped together so that the acoustic condition alternates at the group level (2 sentences) within a trial during presentation. The linguistic content between neighbouring groups could be either matched (same sentences) or unmatched (different sentences) and the relative position of the sentence groups of different acoustic conditions also varied (Figure 1). This produced four different experimental conditions in total. For the two unmatched conditions (“natural speech” precedes or succeeds “8 channel noise vocoding”), the first and second set of 60 sentences were both divided into two equal subsets. From each 30 sentences subset, 6 “natural speech” sentences and 6 “8 channel noise vocoding” sentences were randomly drawn for the first trials, another 6 randomly drawn from the remainder of 24, and so on to produce 5 trials of 12 sentences for each condition. For the two matching conditions, the same set of 60 sentences (third set) was used to generate 60 “8 channel noise vocoding” sentences and then the same operations were applied as for the mismatching conditions. This trial generation process was repeated six times to produce a total of 30 trials for each condition for the whole experiment. Over the 30 trials, each sentence was repeated six times. In each trial, 12 sentences (6 pairs) with alternating acoustic conditions were presented isochronously (We note that this method of stimulus construction and presentation results in speech that is significantly less intelligible than more naturalistic (non-isochronous) speech (Smith et al., 2002; Ding et al., 2014)).

**Figure 1:**
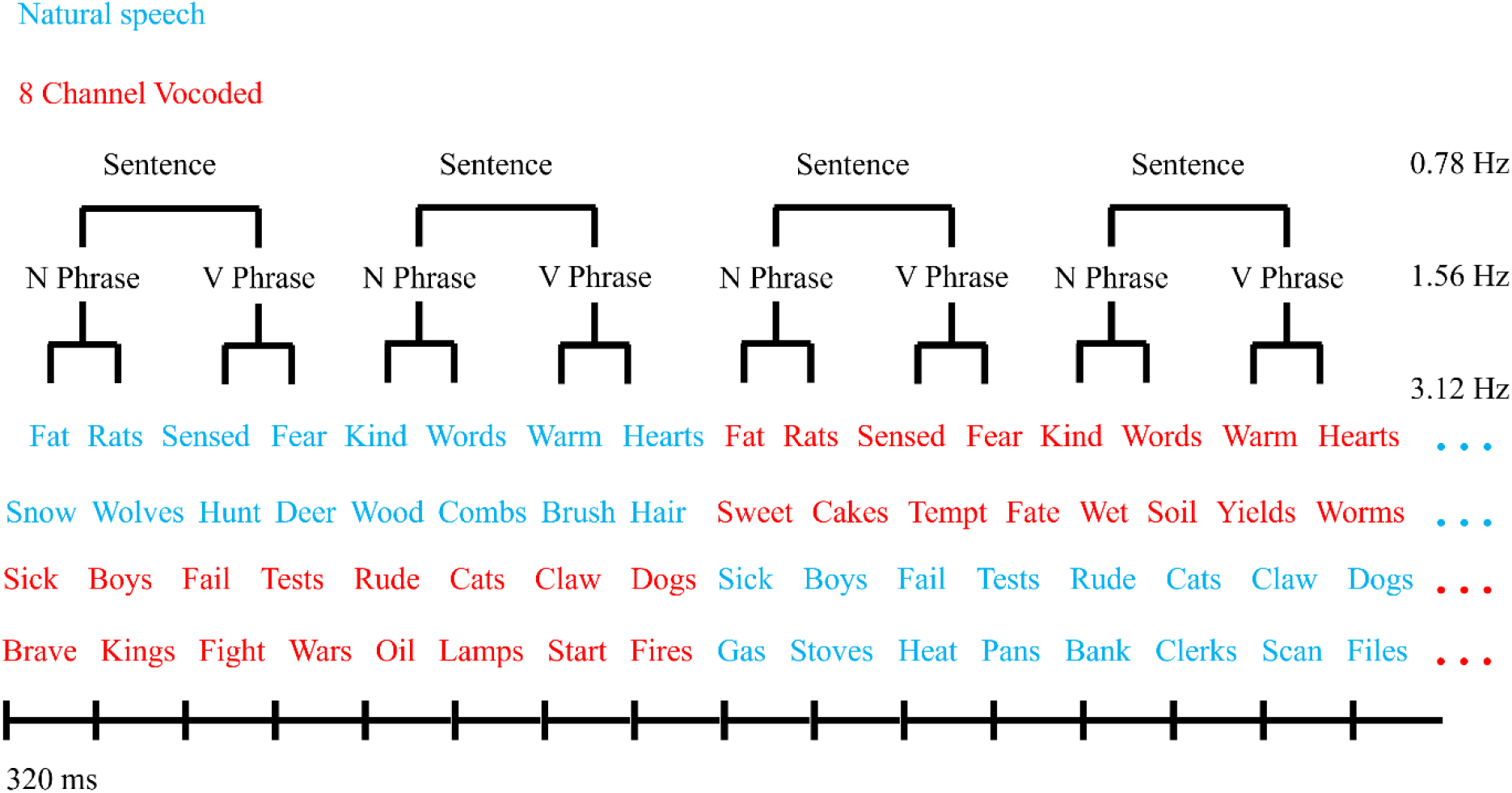
Sequences of English monosyllabic words were presented isochronously, forming phrases and sentences. Acoustic condition alternates between natural and 8 channel noise vocoding after every two sentences with either the same two sentences (matched) or a different pair (unmatched). N and V represent noun and verb, respectively. The far right column indicates the corresponding presentation rate of the linguistic structures.

Out of the 30 trials, there were 6 catch trials constructed for each condition (24 normal trials). In catch trials, 4 consecutive words selected from a random position within a trial, were replaced by four random words to abolish any meaningful sentence structure.

A schematic plot of the linguistic units embedded in the isochronously presented syllable streams is depicted in **Figure 1** below:

#### 2.2.1. Noise Vocoding

In total, 120 four-syllable sentences (including the 60 sentences for “8 channel noise vocoding” and 60 sentences for both acoustic conditions) from the sentence list were processed with noise vocoding to degrade intelligibility. Noise vocoding was implemented using custom MATLAB scripts (MATLAB and Statistics Toolbox Release 2017b, The MathWorks, Inc., Natick, Massachusetts, United States). The frequency range of 200 Hz to 22,050 Hz was divided into 8 logarithmically spaced channels using a 6^th^ order Butterworth filter. In each frequency channel, the envelope of the speech stimulus was extracted with half-wave rectification and a low-pass filtering at 300 Hz (2^nd^ order Butterworth filter). This envelope was then used to amplitude modulate white noise filtered into the same frequency channel from which the envelope was extracted. These envelope-modulated noises were then recombined over frequency channels to yield the noise-vocoded speech segments. The root-mean-square (RMS) level of the noise-vocoded stimulus was normalized to match that of the original speech signal.

To validate and quantify the effect of prior knowledge on intelligibility manipulation, a behavioural word report task was performed by a separate group of native English speakers (n = 30; 18 - 44 years old, mean 21 years old; 21 female). A major part of the participants data (n = 26) was collected online with Gorilla Experiment Builder (Anwyl-Irvine, Massonnié, Flitton, Kirkham, & Evershed, 2020). In this task, participants heard two isochronously presented four-syllable English sentences (one “natural speech” and one “8 channel noise vocoding”) at a time and were required to type only the second sentence out on the screen using a keyboard. The linguistic content (matched / unmatched) and relative position of these two sentences were both varied to produce four different experimental conditions.

Several example trials from each condition were played first to each participant and then sentence pairs were presented in 4 separate blocks (30s sentence pairs/block, all conditions intermixed and evenly distributed) at a comfortable listening level. Participants indicated they had finished typing by pressing the return key, which initiated presentation of the next trial with a delay at 1.2 s. The percentage of correctly reported words is shown in **Figure 2**. As expected, when natural speech precedes vocoded speech, accuracy for the matched condition was significantly higher than for the unmatched condition (mean = 86.9% and 24.5%, *p* < 0.001, paired one-sided t test) while the accuracy for the two conditions did not differ from each other when natural speech succeeds vocoded speech (mean = 87.6% and 87%, *p* = 0.625, paired one-sided t test).

**Figure 2:**
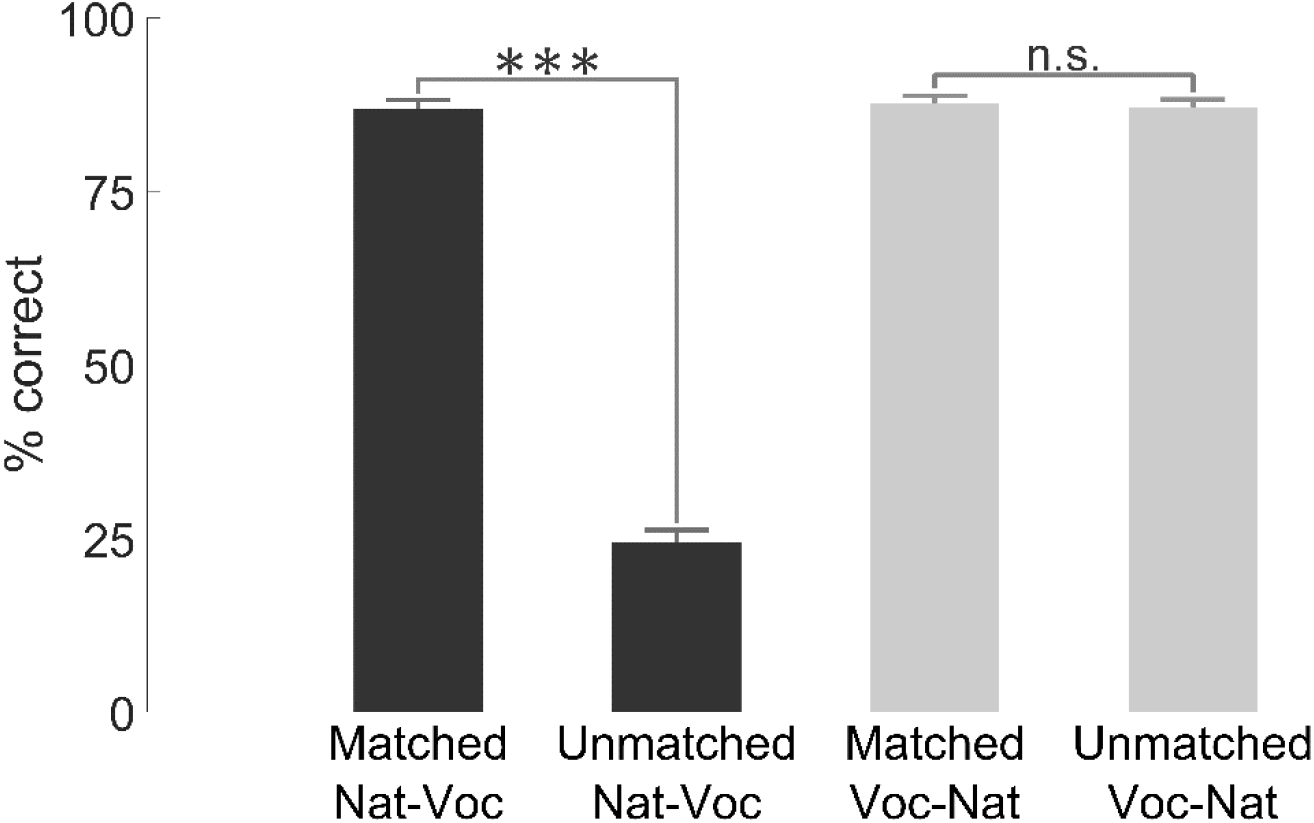
Performance of the word report task under different conditions of prior knowledge (matched speech: natural to vocoded, unmatched speech: natural to vocoded, matched speech: vocoded to natural and unmatched speech: vocoded to natural, error bars reflect standard errors of mean, SEM; the stars indicate the significance levels of 0.001(***), n.s. means not significant).

#### 2.2.2. Stimulus Characterization

The slowly varying temporal envelope of speech signal reflects sound intensity fluctuations and was therefore used to characterize the acoustic property of the stimuli. The amplitude envelope of each trial (12 sentences) was extracted using half-wave rectification followed by a low-pass filtering (cut-off at 30 Hz). The mean power spectrum shown in **Figure 3** was acquired by applying a Fast Fourier Transform (FFT) to individual amplitude envelopes and then averaging within each condition. It is evident that across the four conditions, the envelopes of each stimulus all exhibited strong power modulations at the syllable rate. No such power modulations are observed at phrase or sentence rates.

**Figure 3:**
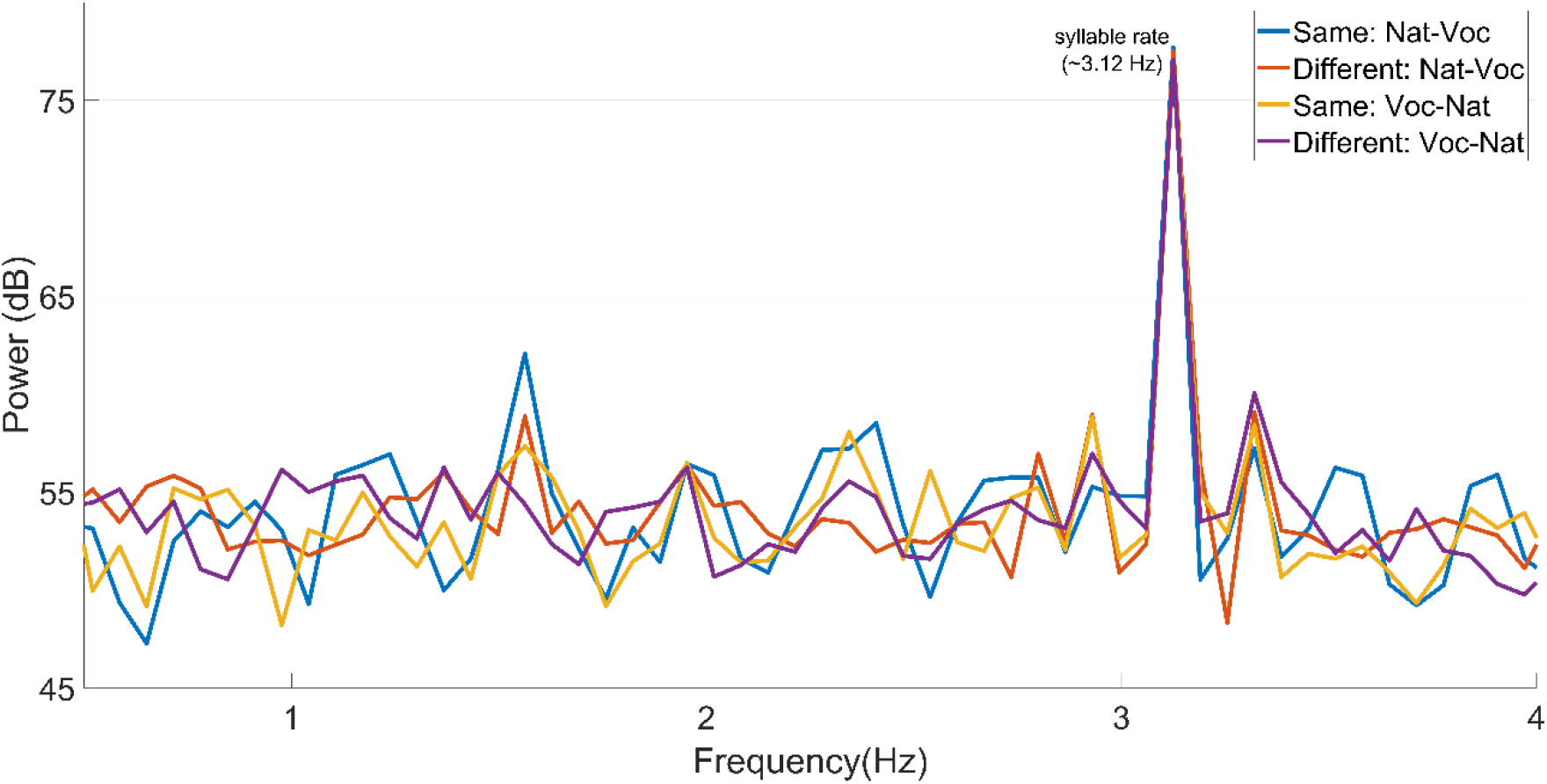
Acoustic characteristics of the speech stimuli. Power spectra of different speech stimuli. Across the four different conditions, the stimulus power was all strongly modulated at the syllable rate (∼3.12 Hz) but not at phrase (∼1.56 Hz) or sentence rates (∼0.78 Hz) (see Figure 1).

### 2.3. Experimental Procedure

Example trials from each condition were played to each participant prior to the experiment. Experiment trials from all conditions were intermixed and evenly distributed into 4 different blocks at 75dB sound pressure level (SPL) through custom built, high fidelity insert earphones (Raicevich, Burwood, Dillon, Johnson, & Crain, 2010) with a flat frequency response up to 8 kHz. Participants were instructed to fix their gaze on a central cross projected to a ceiling screen and indicate whether it was a normal trial or a catch trial (containing at least one ungrammatical sentence) via a button press (with index or middle finger, right hand) at the end of each trial. The button press also initiated presentation of the next trial with a randomly-selected delay of 1.2 s, 1.4 s or 1.6 s. Each block had 24 normal trials and 6 catch trials and the trial types were presented in a pseudo-random order. A technical error resulted in 2 normal trials from each condition incorrectly marked as catch trials for one participant and one normal trial from a single condition (unmatched speech: Voc - Nat) was not recorded for another participant. Analysis was carried out with 22 trials across all conditions and 23 trials for that particular condition for these two participants.

### 2.4. MEG & MRI Data Collection

Prior to MEG recordings, marker coil positions and head shape were measured with a pen digitizer (Polhemus Fastrack, Colchester, VT). Brain activity was recorded continuously using the KIT-Macquarie MEG160 (Model PQ1160R-N2, KIT, Kanazawa, Japan), a whole-head MEG system consisting of 160 first-order axial gradiometers with a 50-mm baseline (Kado et al., 1999; Uehara et al., 2003). MEG data was acquired with the analogue filter settings as 0.03 Hz high-pass, 200 Hz low-pass, power line noise pass through and A/D convertor settings as 1000 Hz sampling rate and 16-bit quantization precision. The measurements were carried out with participants in a supine position in a magnetically shielded room (Fujihara Co. Ltd., Tokyo, Japan). Marker coils positions were also measured before and after each recording block to quantify participants’ head movement, the displacements were all below 5 mm. The total duration of the experiment was about 45 minutes.

Magnetic resonance images (MRI) of the head were acquired for all 23 participants at the Macquarie University Hospital, Sydney, using a 3 Tesla Siemens Magnetom Verio scanner with a 12-channel head coil. Images were acquired using an MP-RAGE sequence (208 axial slices, TR = 2000 ms, TE = 3.94 s, FOV = 240 mm, voxel size= 0.9 mm3, TI = 900, flip angle = 9°).

### 2.5. Data Analysis

MEG data analysis was performed on normal trials only (excluding the catch trials), using the open-source FieldTrip toolbox (Oostenveld, Fries, Maris, & Schoffelen, 2011) and custom MATLAB scripts. Offline MEG data were first filtered with a high-pass filter (0.1 Hz), a low-pass filter (30 Hz) and notch filters (50 Hz, 100 Hz, 150 Hz) and then segmented into epochs according to trial definition. To avoid excessive onset evoked responses, only the data between the start of the second sentence (or the fifth syllable if the stimulus contained no sentential structure) and the end of each trial were analysed further (14.08 s). All data trials were down-sampled to 200 Hz prior to independent component analysis (ICA) (Makeig, Bell, Jung, & Sejnowski, 1996) to remove eye-blinks, eye-movements, heartbeat-related artefacts and magnetic jumps. Components corresponding to those artefacts were identified as by their spectral, topographical and time course characteristics. All cleaned trials of MEG data were kept after ICA artefact rejection.

#### 2.5.1. Sensor Level Analysis

The specific form of this cortical tracking of hierarchical linguistic structure has been demonstrated in a study from Zhang and Ding (Zhang & Ding, 2017) as slow neural fluctuations that emerge at the beginning of the stimulus onset, rather than a series of transient responses at boundaries. Motivated by these characteristics, data analysis was carried out in the frequency domain to reveal brain activities tracking the different levels of linguistic units.

We calculated the magnitude-squared coherence between the MEG recordings and a composite signal following the same analysis procedure described in our recent MEG study (Meng et al., 2021). Magnitude-squared coherence is a frequency-domain measure of phase consistency between two signals across multiple measurements, with a normalized value between 0 and 1 at distinct frequencies. Therefore, phase relationships between these sinewaves in the composite signal can be arbitrary. MEG data trials, as well as the composite signal, were segmented into short frames of 1.28-sec in length and transformed to the frequency domain with FFT and using a sliding Hanning window (75% overlap, 41 frames/trial, ∼ 0.78 Hz frequency resolution). Coherence was then calculated with the power spectral density of each MEG channel and the cross-spectral density between each MEG channel and the composite signal, estimated from the frequency transformed data frames.

#### 2.5.2. Source Analysis

To investigate the spatial distribution of cortical areas coherent to different levels of linguistic structure, we conducted a whole-brain beamforming analysis using Dynamic Imaging of Coherent Sources (DICS) (Gross et al., 2001) which is a frequency domain, linearly constrained minimum variance beamformer (Veen, Drongelen, Yuchtman, & Suzuki, 1997). Source models were constructed based on each participant’s structural MRI. Cortical surface reconstruction (white-grey matter boundary) and volumetric segmentation was performed with the FreeSurfer software suite ((Fischl, 2012); http://surfer.nmr.mgh.harvard.edu/). Cortical mesh decimation (ld factor 10 resulting in 1002 vertices per hemisphere) and surface-based alignment was performed with SUMA - AFNI Surface Mapper (Saad & Reynolds, 2012). A single shell volume conduction model (Nolte, 2003) was adopted and the 2004 cortical surface vertices were used as MEG sources for the leadfield calculation. For more details of the source head modelling procedure, see (Li Hegner et al., 2018).

DICS was applied to the FFT transformed MEG data frames at the corresponding frequency of each linguistic unit across all intelligibility conditions. Coefficients characterizing the beamformer were computed from the cross-spectral density matrix and leadfield matrix at the dominant orientation. Source level coherence images were generated by calculating coherence values between neural activity at each vertex (source point) and the composite signal using the resulting beamformer coefficients. Random coherence images were generated as the average of 100 source space coherence values calculated using the same composite signal but were randomly shuffled at each time, similar to the implementation described by Peelle et al. (2013). Cortical level group analyses were performed using cluster-based permutation test to correct for multiple comparisons (Maris & Oostenveld, 2007) with a critical value of alpha = 0.01 and 2000 random permutations. Each coherence image was contrasted with the corresponding random coherence image; the effect of immediate prior knowledge was evaluated by contrasting coherence images between the matched speech and unmatched speech conditions.

## 3. Results

### 3.1. Behavioural Results

Averaged across all 30 trials under each experimental condition, accuracy rates for the vigilance task (indicate whether sentences were grammatical or not) are calculated as 68.8%, 60.0%, 72.8% and 57.1% respectively (summarised in Table 1). Assessed by paired two-sided t tests, accuracies were significantly higher for the matched (same speech) condition than for the unmatched (different speech) condition, whether natural speech was presented first (p = 0.042) or when vocoded speech was presented first (p < 0.001). This result reflects the greater difficulty of the “different/unmatched speech” conditions.

**Table 1:** Behavioural performance for all experimental conditions (accuracy rate mean ± SEM)

| Matched: Nat-Voc | Unmatched: Nat-Voc | Matched: Voc-Nat | Unmatched: Voc-Nat |
| --- | --- | --- | --- |
| 68.8 ± 2.7% | 60.0 ± 3.8% | 72.8 ± 2.3% | 57.1 ± 2.9% |

### 3.2. Phase-Locked Responses to Hierarchical Linguistic Structures

Magnitude-squared coherence with the composite signal calculated under each condition was grand averaged across all MEG channels as well as all participants and plotted in **Figure 4.** Compared with the averaged power spectra of speech stimuli (**Figure 3**), it is evident that the coherence plot exhibits peaks corresponding to the sentence rate (∼0.78 Hz), phrase rate (∼1.56 Hz) and the syllable rate (∼3.12 Hz). At the sentence and phrase levels, mean coherence values are significantly higher for the matched speech condition than for the unmatched speech condition when natural speech was presented first (sentence level: *t*(22) = 2.61, *p* = 0.016, *r* = 0.49, phrase level: *t*(22) = 3.75, *p* < 0.01, *r* = 0.63, paired two-sided *t* test). In contrast, the mean coherence values at these two levels were not significantly affected by changes in prior knowledge (matched versus unmatched) when noise vocoded speech was presented prior to natural speech (sentence level: *t*(22) = -0.22, *p* = 0.824, phrase level: *t*(22) = -1.05, *p* = 0.31, paired two-sided *t* test). At the syllable level, coherence values were not significantly different between matched and unmatched speech condition in either order of presentation (Nat-Voc: *t*(22) = 1.90, *p* = 0.07, Voc-Nat: *t*(22) = 0.45, *p* = 0.66, paired two-sided *t* test).

**Figure 4:**
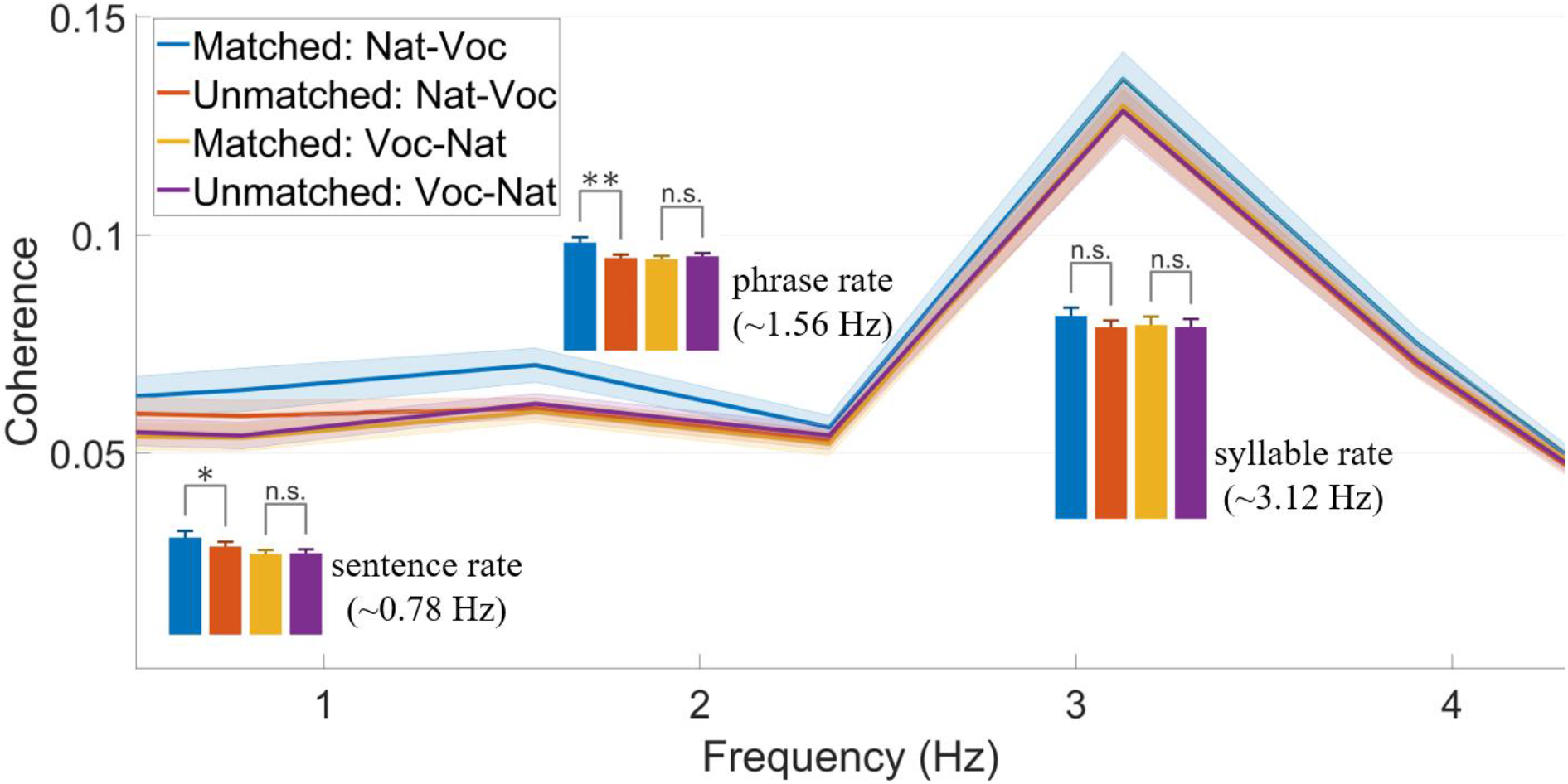
Cortical tracking responses and perceptual context. Averaged MEG sensor level coherence between each MEG channel (160 channels in total) and the composite signal exhibited different tracking activity to the hierarchical linguistic information (syllable, phrase and sentence). The shaded area indicates ± SEM. The bar charts display averaged coherence at distinct frequencies corresponding to hierarchical linguistic units under different conditions (error bars reflect SEM; the stars indicate the significance levels of 0.05(*) and 0.01(**), n.s. means not significant).

### 3.3. Cortical Sources Coherent to Hierarchical Linguistic Structures

The DICS source localization results (quantified as coherence values) were overlaid on the cortical mesh of each individual participant. For visualization purposes, source space results were grand averaged and plotted on a common brain mesh generated using the Freesurfer template brain (http://surfer.nmr.mgh.harvard.edu/), segmented and processed following the procedure described in the Data Analysis section.

**Figure 5** shows grand mean source coherence results for each experimental condition and linguistic unit. Several features are worth noting prior to statistical analyses. First, mean coherence at the syllable level was bilateral and similar in size, in both hemispheres, across all experimental conditions. Second, when the natural speech was presented prior to vocoded speech, mean coherence values at the phrase and sentence levels were larger in the left hemisphere for the matched speech condition than for the unmatched speech condition. Whereas when vocoded speech was presented first, mean coherence values under match and unmatched condition resembled each other at the phrase and sentence levels for both hemispheres.

**Figure 5:**
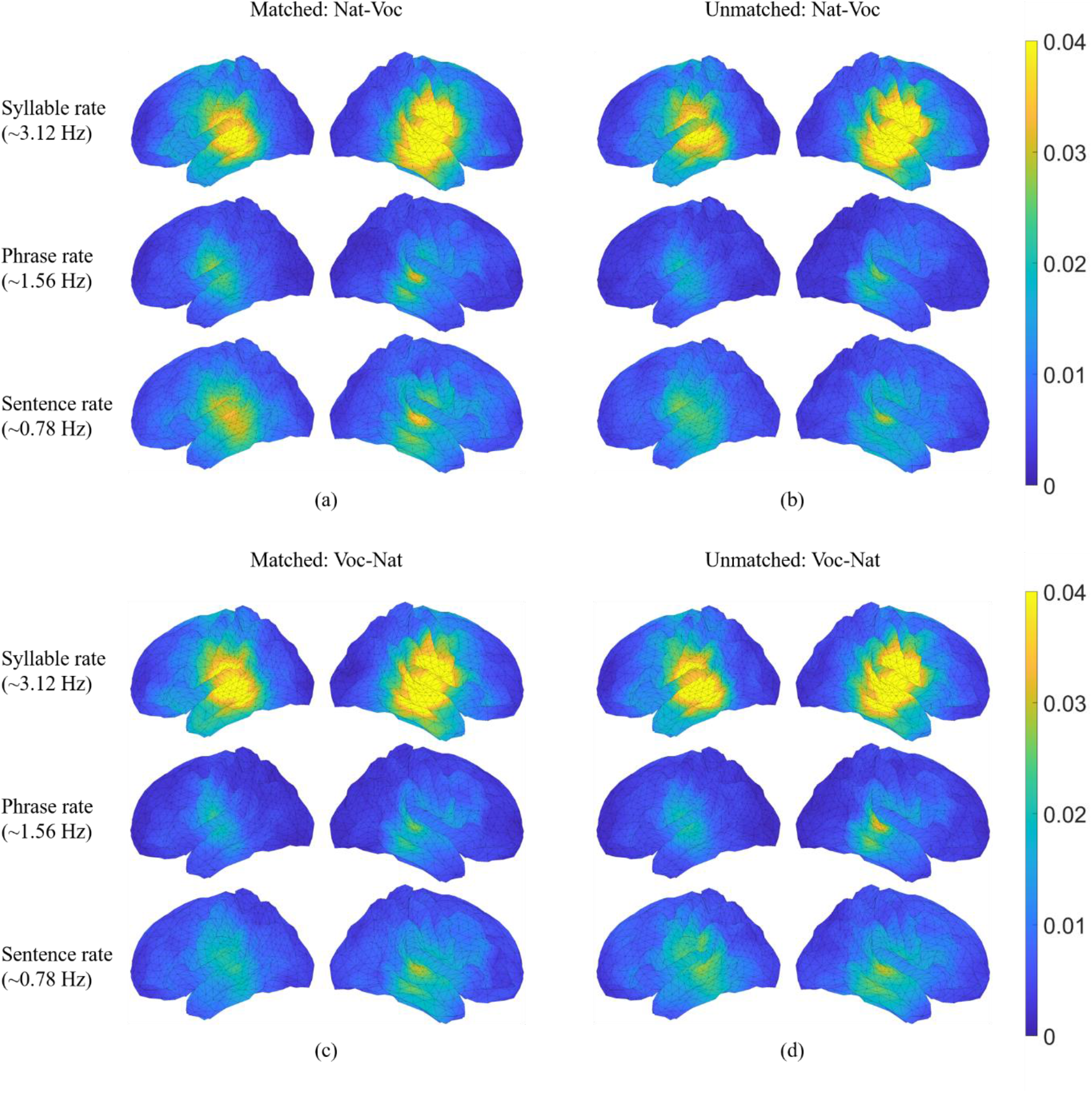
Grand mean source plots. Grand averaged coherence values at frequencies corresponding to syllable rate are sustained across all contextual conditions. (a): Grand averaged coherence values at frequencies corresponding to phrase and sentence rates are left-lateralized when the same natural speech was presented first. (b): Grand averaged coherence values at frequencies corresponding to phrase and sentence rates are reduced when the prior natural speech was different and do not show clear lateralization. (c) and (d): Grand averaged coherence values at frequencies corresponding to phrase and sentence rates are bilateral and do not show clear difference between matched and unmatched conditions when vocoded speech was presented first. Colour bars indicate coherence values.

#### 3.3.1. Contrast with random coherence

Whole-brain analyses contrasted coherence maps in each experimental condition against “random” coherence maps (calculated using shuffled composite signals – see Methods section). Results are shown in **Figure 6** using paired one-sided t tests with a sample-wise threshold of *p* < 0.0005 and a threshold of *p* < 0.0005 whole-brain cluster extent multiple comparison correction.

**Figure 6:**
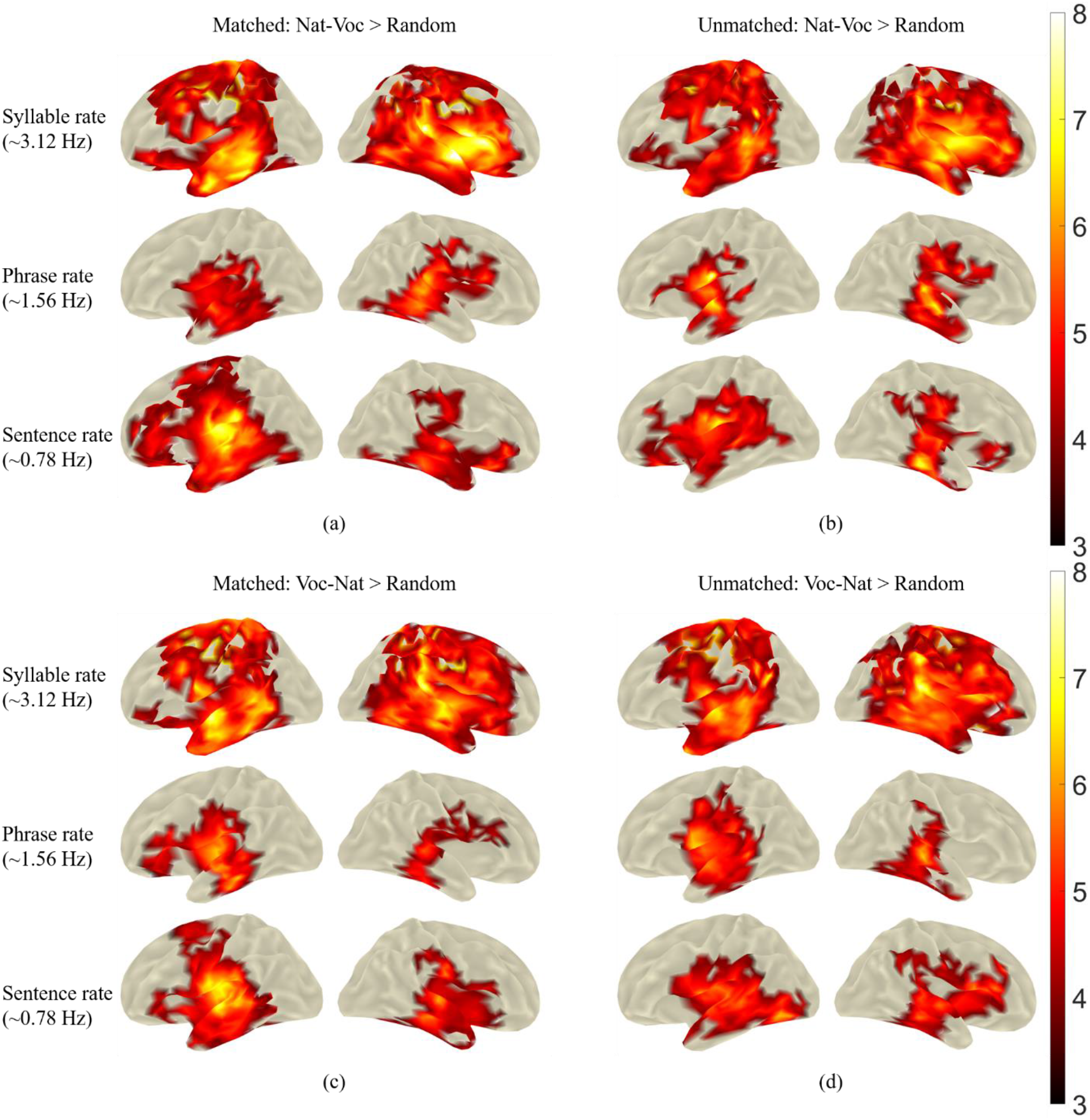
Contrasts with random coherence. Bilateral coherence is observed at all three frequencies. (a) Matched speech, natural speech presented prior to vocoded speech. (b) Unmatched speech, natural speech presented prior to vocoded speech. (c) Matched speech, vocoded speech presented prior to natural speech. (d) Unmatched speech, vocoded speech presented prior to natural speech. Colour bar indicates t values.

The results show that significantly higher coherence to all levels of linguistic structure in bilateral peri-Sylvian cortices across all contextual conditions (relative to coherence to randomly shuffled composite signals).

#### 3.3.2. Contrasts against shuffled speech

The foregoing contrasts provide a picture of the overall extent to which our measured neuronal responses tracked each of the three rates in the composite signal. To examine the effect brought about by immediate prior experience with the acoustic and linguistic information on cortical tracking activity, we conducted a whole-brain search for regions in which the coherence value was higher for the matched speech condition than the unmatched speech condition when the natural speech was presented either prior to or after the noise vocoded speech.

As shown in **Figure 7**, positive clusters were found when the same natural speech was presented before noise vocoded speech using a vertex-wise threshold of p < 0.05 and whole-brain cluster extent correction for multiple comparison at p < 0.01. The enhancements in coherence were restricted to the left hemisphere at phrase and sentence level. Notably, no significant clusters were obtained for the syllable rate contrast (top row) or for the contrast between the matched condition and unmatched condition when the noise vocoded speech was presented prior to the natural speech.

**Figure 7:**
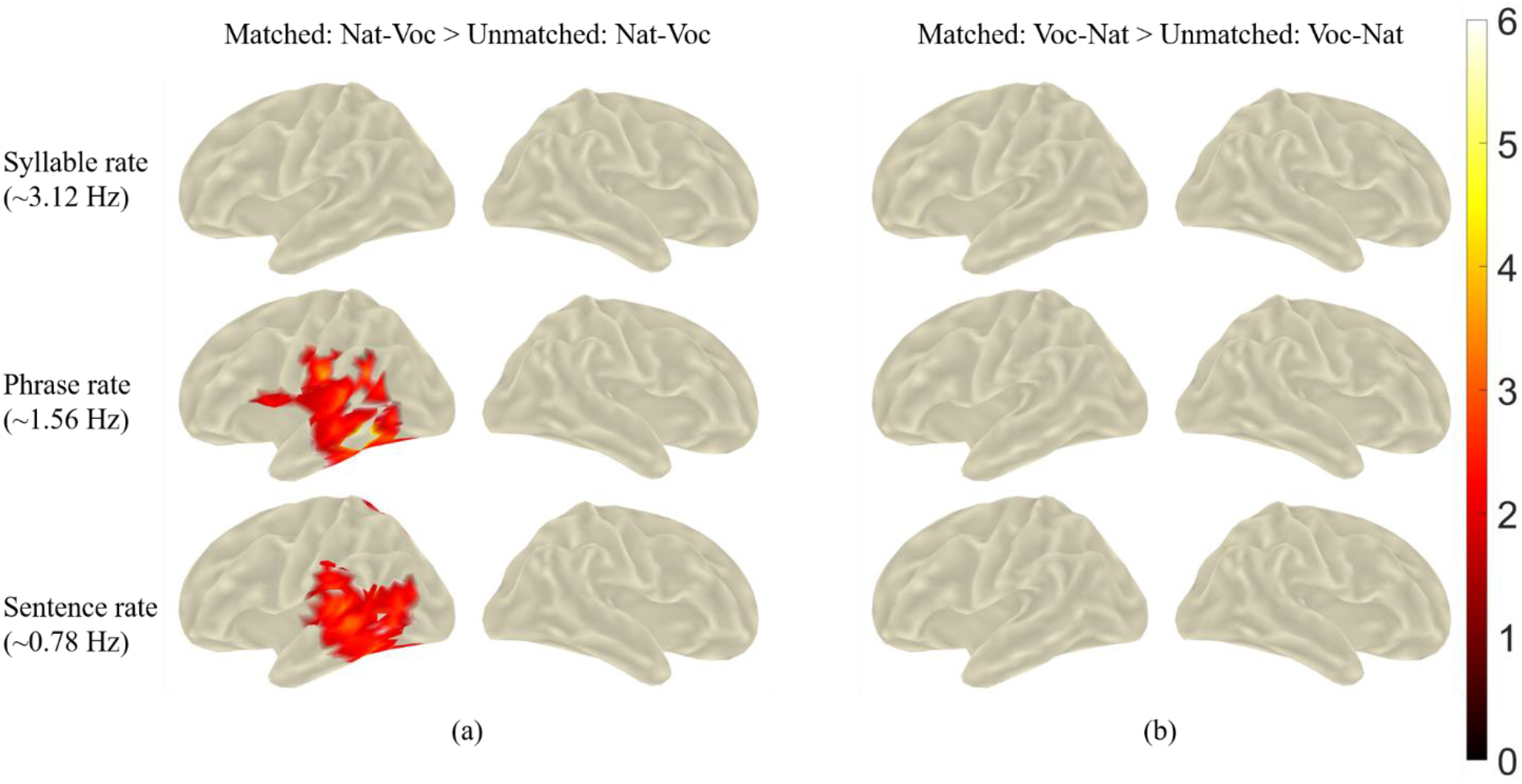
Contrasts against unmatched speech. (a) Cortical regions showing enhanced coherence under matched speech condition compared to the unmatched speech condition when natural speech was presented first. (b) No significant clusters were found for the contrast between matched condition and unmatched condition when noise vocoded speech was presented prior to natural speech. Colour bar indicates t values.

## 4. Discussion

A compelling neurophysiological basis for the well-studied perceptual “pop-out” effect has recently been provided from electrophysiological data recorded directly from the human auditory cortex (Holdgraf et al., 2016): Prior exposure to intact speech results in rapid and automatic enhancements in neural representations of key spectrotemporal features which can be employed during the subsequent identification of these signals in degraded form. However experience-driven neurophysiological effects have not been convincingly demonstrated in several studies using measures of neural entrainment (Holdgraf et al., 2016; Liberto et al., 2018; Millman et al., 2014), which are proposed to be necessary for speech comprehension. The results of the current study show that when speech intelligibility was enhanced by matched prior acoustic and linguistic knowledge, a corresponding enhancement can be obtained in MEG responses to embedded linguistic units, effects that were evident at the phrase and sentence level. Statistical analyses at the source level revealed regions of greater coherence in the left temporal cortex to phrase and sentence level regularities. When prior information was abolished by presenting the noise-vocoded speech prior to the natural speech, no perceptual pop-out was achieved and there were also no differences in the corresponding MEG responses.

Our finding of no significant effect at the syllable level is consistent with the results of previous investigations of the perceptual “pop-out” effect (Holdgraf et al., 2016; Liberto et al., 2018; Millman et al., 2014), showing no effects on the cortical phase-locking response to the speech envelope (which reflects modulations at the syllable rate). As discussed in our previous MEG study (Meng et al., 2021) and the EEG study from Liberto et al. (2018), the lack of effect on the syllable-level responses is likely to be attributable to the fact that this response is driven largely by physical regularities present in the speech stream and is consequently less susceptible to top-down influences induced by prior knowledge.

The ubiquitous oscillatory neural activities in the brain have been proposed to provide a potential brain mechanism for deciphering the speech signal (Giraud & Poeppel, 2012; Hickok & Poeppel, 2007). Many neuroimaging studies have examined the low frequency entrainment to slow varying speech envelope in auditory cortex, as the putative brain process segregating linguistic units at syllable scale. However the results to date have been mixed and controversial (Ding & Simon, 2014; Peelle & Davis, 2012; Zoefel & VanRullen, 2015). The neural response to hierarchical linguistic structures (Ding et al., 2016) provides a plausible mechanism for information integration over time (Buzsáki, 2010; Schroeder, Lakatos, Kajikawa, Partan, & Puce, 2008) and enables structure building operations (Bastiaansen, Magyari, & Hagoort, 2009) via coupling with higher frequency neural oscillations (Canolty et al., 2006; Lakatos et al., 2005; Sirota, Csicsvari, Buhl, & Buzsáki, 2003). Therefore, examining this hierarchy of neural processing may provide insights into the delineating process of those controversial results from speech envelope tracking measurement, e.g., the perceptual “pop-out” effect facilitated by prior knowledge and top-down integration.

Our statistical maps indicate that the experience-dependent enhancement of the tracking of phrase- and sentence-level responses are associated with activity in the left cerebral hemisphere. The lateralised responses are entirely consistent with the findings reported in our recent MEG study demonstrating that the speech intelligibility modulates changes in cortical tracking responses to larger linguistic units (phrases and sentences) (Meng et al., 2021).

The issue of hemispheric specialisations for speech analysis is complex (Poeppel, Emmorey, Hickok, & Pylkkänen, 2012, p. 201) and strongly debated. Previous neuroimaging studies of the perceptual “pop-out” phenomenon have reported left hemisphere activation (Dehaene-Lambertz et al., 2005; Di Liberto et al., 2018), bilateral activation (Giraud et al., 2004; Liebenthal et al., 2003; Sohoglu & Davis, 2016; Sohoglu et al., 2012; Tuennerhoff & Noppeney, 2016), or no effect (Millman et al., 2014). Result of the present study also indicate that these contradictions can be partly or largely attributed to the confound between acoustic and linguistic cues in the speech stimuli employed in these studies. In contrast, when physical and linguistic cues are unambiguously dissociated, our results showed a clear left hemisphere lateralization of cortical sources coherent to phrase and sentence level linguistic regularities, after the perceptual “pop-out” occurred. There has been an emerging consensus from researchers using fMRI to measure hemodynamic responses to acoustic and speech stimuli. Based on the results from a series of fMRI studies of speech comprehension, Peelle (2012) concluded that cortical lateralisation depends in a roughly graded fashion on the relative amounts of linguistic processing required by the task. With a single experiment showing concurrent responses to different levels of linguistic units, our current MEG results provide a clear confirmation of this conclusion.

Since the prior knowledge driven enhancement in speech intelligibility relies on linguistic attributes, the cortical origins of this rapid tuning shift has been predicted to be within the auditory association areas or the non-auditory regions such as the inferior frontal gyrus/premotor cortices (Holdgraf et al., 2016). In support of this prediction, our source analyses point to frontal-temporal origins of the top-down processes for the brain tracking responses at phrase and sentence level, mainly encompassing ventral motor and premotor regions; and not to more anterior pre-frontal executive regions. Contrary evidence has been reported in an earlier MEG study, suggesting that an early activation of left IFG initiates subsequent early speech envelope entrainment activities in left HG, STS and MTG (Di Liberto et al., 2018). It has been demonstrated by Meng et al. (2021) that brain responses to mixed acoustic and linguistic cues are largely driven by the acoustic cues. For studies using naturalistic speech stimuli to investigate the brain mechanism of speech envelope encoding/speech envelope entrainment, such as the one by Di Liberto and colleagues, such a confound between acoustic and linguistic cues is inevitable. For the current study, we measured and reported brain responses that are unambiguously dissociated from any acoustic cues and this may explain the discrepancies with prior literature.

As our experiment results demonstrated, the MEG responses exhibit a fast plasticity driven by prior knowledge with intelligible speech. This neural tracking activity nicely characterizes the dominant factors involved in linguistic processing, from both the bottom-up and top-down process perspectives. It therefore serves well as an objective neural marker of high-level speech processing which will be useful in basic neurolinguistics research, and also has potential clinical significance for assessment of language function after interventions for hearing loss including cochlear implantation. A growing consensus in the field of cochlear implantation is that much of the observed variability in performance may be attributable to neuroplastic changes in speech processing as a consequence of the profound sensory deprivation imposed by deafness (Wilson & Dorman, 2008). The present results, taken together with the successful measurement of brain tracking responses to hierarchical linguistic units using electroencephalography (EEG) by Ding et al. (2017), provides a potentially powerful neuromarker that can be used to assess and interrogate the “compromised auditory brains” of cochlear implant (CI) recipients after the restoration of auditory inputs. Objective markers of language processing may also be useful in studies of young children and in difficult-to-test clinical populations including autism spectrum disorders.

## Supporting information

Supplementary Material: Sentence Stimuli

## 5. Author Contributions

Q.M., Y.L.H., C.M and B.J. conceived and designed the experiment. Q.M. performed the MEG experiments. QM and Y.L.H performed data analysis. QM, Y.L.H and B.J wrote the paper. Y.L.H. and B.J. contributed equally as senior authors. All of the authors discussed the results and edited the manuscript.

